# Hybridization capture of larch (*Larix* Mill) chloroplast genomes from sedimentary ancient DNA reveals past changes of Siberian forests

**DOI:** 10.1101/2020.01.06.896068

**Authors:** Luise Schulte, Nadine Bernhardt, Kathleen R. Stoof-Leichsenring, Heike H. Zimmermann, Luidmila A. Pestryakova, Laura S. Epp, Ulrike Herzschuh

## Abstract

Siberian larch (*Larix* Mill.) forests dominate vast areas of northern Russia and contribute important ecosystem services to the world. It is important to understand the past dynamics of larches, in order to predict their likely response to a changing climate in the future. Sedimentary ancient DNA extracted from lake sediment cores can serve as archives to study past vegetation. However, the traditional method of studying sedimentary ancient DNA – metabarcoding – focuses on small fragments which cannot resolve *Larix* to species level nor allow the detailed study of population dynamics. Here we use shotgun sequencing and hybridization capture with long-range PCR-generated baits covering the complete *Larix* chloroplast genome to study *Larix* populations from a sediment core reaching back up to 6700 years in age from the Taymyr region in northern Siberia. In comparison to shotgun sequencing, hybridization capture results in an increase of taxonomically classified reads by several orders of magnitude and the recovery of near-complete chloroplast genomes of *Larix*. Variation in the chloroplast reads corroborate an invasion of *Larix gmelinii* into the range of *Larix sibirica* before 6700 years ago. Since then, both species have been present at the site, although larch populations have decreased with only a few trees remaining in what was once a forested area. This study demonstrates for the first time that hybridization capture applied to ancient DNA from lake sediments can provide genome-scale information and is a viable tool for studying past changes of a specific taxon.

## Introduction

Siberian forests are unique as they cover a vast area of about 263.2 million ha (Abaimov, 2010) dominated by a single genus of tree, the deciduous conifer larch (*Larix* Mill.). As the only extensive forest biome growing on continuous permafrost, it plays an important role for local communities and it provides critical ecosystem services in a global context including carbon stocks, climate feedbacks, permafrost stability, biodiversity and economic benefits (Herzschuh, 2019). It is therefore important to understand how the genus and individual larch species will respond to a warming climate.

Frequent natural hybridization between the larch species make it difficult to distinguish taxa and the number of accepted species is still under discussion (Abaimov, 2010). This is one of the reasons why there is still little known about the population dynamics of Siberian larch species and the question remains of whether there have been migrations of larches in the current postglacial.

Sedimentary ancient DNA (*sed*aDNA) from lakes can act as an archive of the past and has been demonstrated to be a valuable tool in the study of past vegetation history (Jørgensen et al., 2012; Parducci et al., 2017; Willerslev et al., 2003). Most *sed*aDNA studies focus on organellar DNA, as the higher copy number of organelles per cell compared to the nucleus allow a higher chance of preservation. The metabarcoding approach (Taberlet, Coissac, Hajibabaei, & Rieseberg, 2012) applied to DNA extracted from sediments is the most common, robust and fast technique to study past vegetation (Alsos et al., 2018; Niemeyer, Epp, Stoof-Leichsenring, Pestryakova, & Herzschuh, 2017; Pansu et al., 2015). A very short, but highly variable DNA fragment is PCR-amplified out of the pool of DNA fragments and subsequently sequenced using high-throughput sequencing. However, the method is not suited to resolve population dynamics of single species, as metabarcoding markers used for ancient degraded samples must be very short while at the same time flanked by primers that are conserved across a larger taxonomic group. Therefore, their taxonomic resolution is, in most cases, insufficient to resolve closely related species (Taberlet et al. 2007), let alone show sub-specific variation.

Sequencing of the entire DNA extracted from ancient sediments, termed metagenomic shotgun sequencing, has been shown to provide information on the entire taxonomic composition of the sample (e.g., fungi, bacteria, archaea; Ahmed et al. 2018; Parducci et al. 2019). By sequencing complete DNA molecules, it is possible to authenticate ancient sequences versus modern contaminants by their specific *post-mortem* DNA damage patterns towards the ends of the molecules (Ginolhac, Rasmussen, Gilbert, Willerslev, & Orlando, 2011). As it is not restricted to a specific DNA fragment, it also allows the retrieval of many different loci belonging to single species provided they are sufficiently concentrated in the sample. Another advantage is that this method avoids bias introduced by PCR. A major drawback, however, is the immense sequencing effort that must be expended to achieve a sufficient overview of the DNA present in a sample. Most of the sequences retrieved from ancient environmental samples are not assignable to a specific taxon because available sequence databases are still limited, and most assigned sequences are not of eukaryotic origin (Ahmed et al., 2018; Pedersen et al., 2016). Especially in the case of DNA extracted from lake sediments, the ratio of sequences assigned to terrestrial plants to total DNA sequenced is expected to be extremely low (Parducci et al., 2019).

A way to overcome the limitations of shotgun sequencing is to enrich the DNA of the focal species in the samples via hybridization capture prior to sequencing. Here, short fragments of DNA of the species and target sites of interest are used as baits, to which the corresponding sites of interest in ancient DNA libraries are hybridized. This technique, originally developed for modern DNA has already been successfully applied to various ancient samples ranging from single specimens (Ávila-Arcos et al., 2011; Maricic, Whitten, & Pääbo, 2010) to cave sediments (Slon et al., 2017) and permafrost samples (Murchie et al., 2019). To date, it has not been applied to DNA extracted from ancient lake sediments, which are especially challenging due to the high diversity of organisms living in the water and sediments and around the lakes. With the exception of metabarcoding analysis, most ancient DNA studies focus on mammals, mostly using mitochondrial DNA (Carpenter et al., 2013; Dabney et al., 2013; Enk et al., 2016). Plants have received limited attention in ancient DNA research (Parducci et al., 2017), and complete chloroplasts have not yet been targeted for hybridization capture of ancient DNA.

Here we apply shotgun sequencing and a hybridization capture approach to *seda*DNA samples from a small lake in the Taymyr region of north-eastern Siberia. The study site lies in the boundary zone of two larch species, *Larix gmelinii* and *Larix sibirica* (Abaimov, 2010), with hybridization occurring between the boundary populations (Abaimov, 2010; Polezhaeva, Lascoux, & Semerikov, 2010). It has been hypothesized for this region, that a natural invasion of *L. gmelinii* into the range of *L. sibirica* occurred during the Holocene (Semerikov, Semerikova, Polezhaeva, Kosintsev, & Lascoux, 2013). The lake is situated in the treeline ecotone with scattered patches of *L. gmelinii* occurring in the area (Klemm, Herzschuh, & Pestryakova, 2016). A sediment core of the lake has already been extensively studied using pollen analysis, DNA metabarcoding and mitochondrial variants (Epp et al., 2018; Klemm et al., 2016) making it an ideal site to study ancient larch population dynamics based on chloroplast DNA.

As a proof of concept, four samples were both shotgun sequenced and enriched for the chloroplast genome of *L. gmelinii*. We compare the proportion of classified reads at the domain level and the coverage of the *Larix* chloroplast genome of these methods. This study highlights the first successful recovery of near-complete chloroplast genomes from ancient lake sediments.

## Methods

### Sample material

Samples were obtained from a sediment core from lake CH12 (72.399°N, 102.289°E, 60 m a.s.l.) in the Khatanga region of the northern Siberian lowlands, located between the Taymyr Peninsula to the north and the Putorana Plateau to the south. The lake’s position is in the northern part of the treeline ecotone and is currently surrounded by a vegetation of single-tree tundra. Details of the chronology of the core are described in Klemm et al. (2016). Four samples were chosen for the present study at depths/ages 121.5 cm/∼6700 calibrated years before present (cal-BP), 87.5 cm/∼5400 cal-BP, 46.5 cm/∼ 1900 cal-BP and 2.5 cm/∼60 cal-BP.

### Laboratory work

#### Sampling, DNA extraction and library preparation

Core sub-sampling and DNA extraction were performed as described in Epp et al. (2018). Five µL of each DNA extraction were used in the library preparation. The extractions had DNA concentrations of 2.4 ng/µL (6700 cal-BP), 5.7 ng/µL (5400 cal-BP), 3.4 ng/µL (1900 cal-BP) and 6.7 ng/µL (60 cal-BP). In total, six libraries with volumes of 50 µL each were prepared: four samples from the CH12 core, one extraction blank and one library blank. Libraries were prepared following the single stranded DNA library preparation protocol of Gansauge et al. (2017) with the following adjustment: the ligation of the second adapter (CL53/CL73) took place in a rotating incubator. The libraries were quantified with qPCR as described by Gansauge and Meyer (2013). The prepared libraries were used downstream for both shotgun sequencing and hybridization capture of chloroplast genomes.

#### Shotgun sequencing

Twenty-four µL of the prepared DNA library were amplified and indexed by PCR with 13 cycles as described in Gansauge and Meyer (2013) using the index primer sequences P5_1 – P5_6 and P7_91 – P7_96. The libraries were pooled in equimolar ratios to a final pool of 10 nM with the two blanks accounting for a molarity of 20% compared to the samples. The sequencing of the pool was performed by Fasteris SA sequencing service (Geneva, Switzerland) using a modified sequencing primer CL72 as described in Gansauge and Meyer (2013). The pool was sequenced on one lane of an Illumina HiSeq 2500 platform (2 x 125 base pairs (bp), High-Output V4 mode).

#### Bait construction

Eighteen primer pairs for long range PCR products covering the whole chloroplast genome of *L. gmelinii* were designed as published in Zimmermann et al. (2019). Long-range PCR products were generated from DNA extractions from needles of a *L. gmelinii* individual collected in the Botanical Gardens of the University of Potsdam (Accession number XX-0POTSD-3867). PCR amplification was conducted using the SequalPrep™ Long PCR Kit (Invitrogen), according to the manufacturer’s cycling protocol instructions, and with specific annealing temperatures for each primer pair (see Zimmermann et al. 2019, Table S2). The PCR products were pooled in equimolar ratios in a volume of 130 µL and sonicated using a Covaris M220 Focused-ultrasonicator (Covaris, Woburn, MA) to a target peak of 350 bp with settings of peak incident power 50 W, duty factor 20%, cycles per burst 200, treatment time 70 seconds. The fragment size and distribution were visualized with Agilent TapeStation (D1000 ScreenTape, Agilent Technologies, Santa Clara, CA). Fragment sizes ranged from 100 bp to 1000 bp with an average size of 370 bp. The complete sonicated pool was purified using the MinElute PCR Purification Kit (Qiagen, Hilden, Germany), following the manufacturer’s recommendations and eluted in 30 µL.

#### Hybridization capture

Another 24 µL of prepared DNA libraries was PCR amplified with 16 cycles and equimolarly pooled with the blanks having a molarity of 20% compared to the samples. The enrichment was done according to the protocol of Maricic, Whitten, and Pääbo (2010b) with the following adjustments: the purified library pool was blunt ended using Fast DNA End Repair Kit (Thermo Scientific) in a volume of 100 µL according to the manufacturer’s recommendations. The adapters were ligated to the baits using the Rapid DNA Ligation Kit (Thermo Scientific) with the following conditions: 15 µL blunt-ended DNA, 1 µL of adapters Bio-T/B (50 µM), 8 µL Quick ligase buffer, 12 µL water and 4 µL quick ligase were mixed and incubated for 15 min at room temperature. In addition to the blocking oligonucleotides provided in the original protocol (Maricic et al., 2010), we used specific oligonucleotides to block the indices used in the double indexed library : BO7.P7.part2.F: ATCTCGTATGCCGTCTTCTGCTTG-phosphate and BO8.P7.part2.R: CAAGCAGAAGACGGCATACGA. Four µL of the final elution were PCR amplified with 18 cycles using the TruSeq DNA Nano preparation kit (Illumina Inc., San Diego, CA), performed by Fasteris SA sequencing service (Geneva, Switzerland). The enriched library pool was sequenced by Fasteris SA sequencing service (Geneva, Switzerland) in the same way as described for shotgun sequencing.

### Data analysis

#### Quality control, trimming and merging of reads

Demultiplexed FASTQ files, as obtained from the sequencing provider, were quality checked using FASTQC (v. 0.7.11, Andrews, 2015) before and after trimming with Trimmomatic (v. 0.35, Bolger, Lohse, & Usadel, 2014). The analyses performed by Trimmomatic relied on a file containing the applied and common Illumina adapters, and the following parameter settings were used: remove adapters with max. mismatch rate = 2, sliding window, window size = 4, average quality = 15, minimum quality to keep a base = 3, minimum length to keep a sequence = 36 nt. Unpaired reads were discarded. Paired reads were merged using PEAR (v. 0.9.10, Zhang, Kobert, Flouri, & Stamatakis, 2014). Merged and unmerged reads were treated in the following steps separately and merged at the end into BAM files with samtools using the command “merge” (v. 1.9, H. Li et al., 2009).

#### Taxonomical classification

Reads were classified using kraken2, (v. 2.0.7-beta, Wood, Lu, & Langmead, 2019) with a conservative confidence threshold (--confidence 0.9) against the non-redundant nucleotide database (nt) from NCBI (ftp://ftp.ncbi.nlm.nih.gov/blast/db/FASTA/nt.gz; downloaded in May 2019) and the NCBI taxonomy (retrieved via the kraken2-build command in June 2019). This classification was used for the description and comparison of the shotgun as well as the capture dataset. Additionally, the reads were classified using a custom database of 8,132 complete plant chloroplast genomes (downloaded from NCBI in July 2019). In this second classification kraken2 was run with default parameters to allow for the retrieval of variation. Reads classified with the chloroplast database as genus *Larix* or below were extracted using custom bash scripts and seqtk (Seqtk, n.d.) and used for further analysis.

#### Alignment

Capture dataset reads classified as *Larix* by comparison to a custom chloroplast database were mapped against a *L. gmelinii* reference genome from an individual in the Taymyr region (NCBI accession no.: MK468637.1). Reads were mapped using BWA aln algorithm (v. 0.7.17-r1188, Li & Durbin, 2009) with the settings -l 1024 -o 2 -n 0.001 to ensure relaxed mapping. Duplicate reads were removed using Picard MarkDuplicates (v. 2.20.2-SNAPSHOT, Broad Institute, 2019) for merged reads and samtools markdup for unmerged reads. The same alignment parameters were used to map the complete, taxonomically unclassified samples to the same *L. gmelinii* chloroplast reference genome.

#### *Coverage of the* Larix *chloroplast genome at different annotated functions*

The annotation of the chloroplast genome was adopted from the published annotation file on NCBI for the used reference genome (Acc.: MK468637.1). Capture dataset alignments of the *Larix*-classified reads and the complete samples against the *Larix* chloroplast genome were used for the analysis of different functional groups. Coverages for each sample were obtained using samtools depth (option –a) (Li et al., 2009) and summed up at each position across the samples. Boxplots were made using R with ggplot2 (R Core Team, 2013).

#### Assessment of ancient DNA damage patterns

Ancient damage patterns were assessed using mapDamage (v. 2.0.8, Jónsson, Ginolhac, Schubert, Johnson, & Orlando, 2013) rescaling the quality scores of the BAM files with the single-stranded option. For damage pattern assessment of unmerged paired-end reads, the reads were mapped in two rounds as single-end reads to the reference (Acc.: MK468637.1) and analysed with mapDamage to not confound the 5’ and 3’ damage patterns. Read length distribution were assessed with mapDamage for overlapping merged reads and with Geneious (v. 2019.2, Biomatters, 2019).

#### Assignment of Larix bases to L. sibirica or L. gmelinii

To distinguish between the two *Larix* species *L. gmelinii* and *L. sibirica* and determine their temporal occurrence, we performed multiple comparisons of the 13 complete chloroplast genomes of *L. gmelinii* (Accessions MK468630–39, MK468646 and -48, NC_044421) available on NCBI and one of the two available *L. sibirica* genomes (Accession: MF795085.1) and classified species-specific single nucleotide polymorphisms (SNPs) and insertions and deletions (indels). The alignment was performed using bwa mem with default parameters. SNPs and indels were called using bcftools mpileup (option -B) and call (option -mv) (v. 1.9, H. Li et al., 2009). In total, 294 positions were determined as occurring only in *L. sibirica*. For each of these positions the above-produced alignments (ancient sample reads against *L. gmelinii* reference) were analysed with regards to whether the reads carried the same variant as *L. gmelinii* or *L. sibirica* or if they had a different SNP or indel at this position. The analysis was carried out using bam-readcount (The McDonnell Genome Institute, 2018) in conjunction with a custom python script which for every position classified the count of reads as *L. gmelinii, L. sibirica* or “other”.

## Results

### Overview of the shotgun and hybridization capture data sets

Shotgun sequencing of the entire DNA extracted from the ancient lake sediment samples resulted in about 424 million read pairs for the four samples and 25 million reads for the two blanks (library and extraction blank). Per-sample read count ranged from 78–145 million paired reads. After trimming and filtering, 62.6% of the reads remained for the analysis, i.e. 265 million (M) reads in total, and 0.1–0.2% for the blanks, i.e. a total of 39 thousand (k) reads. Eighty-two% of the sample reads overlapped and were merged.

Using kraken2 with the nt database, 0.3% of the quality filtered reads could be classified (Figure 1). Bacteria comprise 62.6% of the classified reads (506 k reads across the four samples) and 23.4% were classified as Eukaryota (189 k reads). Of the Eukaryota, the majority was assigned to Viridiplantae (122 k reads) and 2.8 k reads were assigned to *Larix* (Supplementary Table S1).

**Figure 1.**
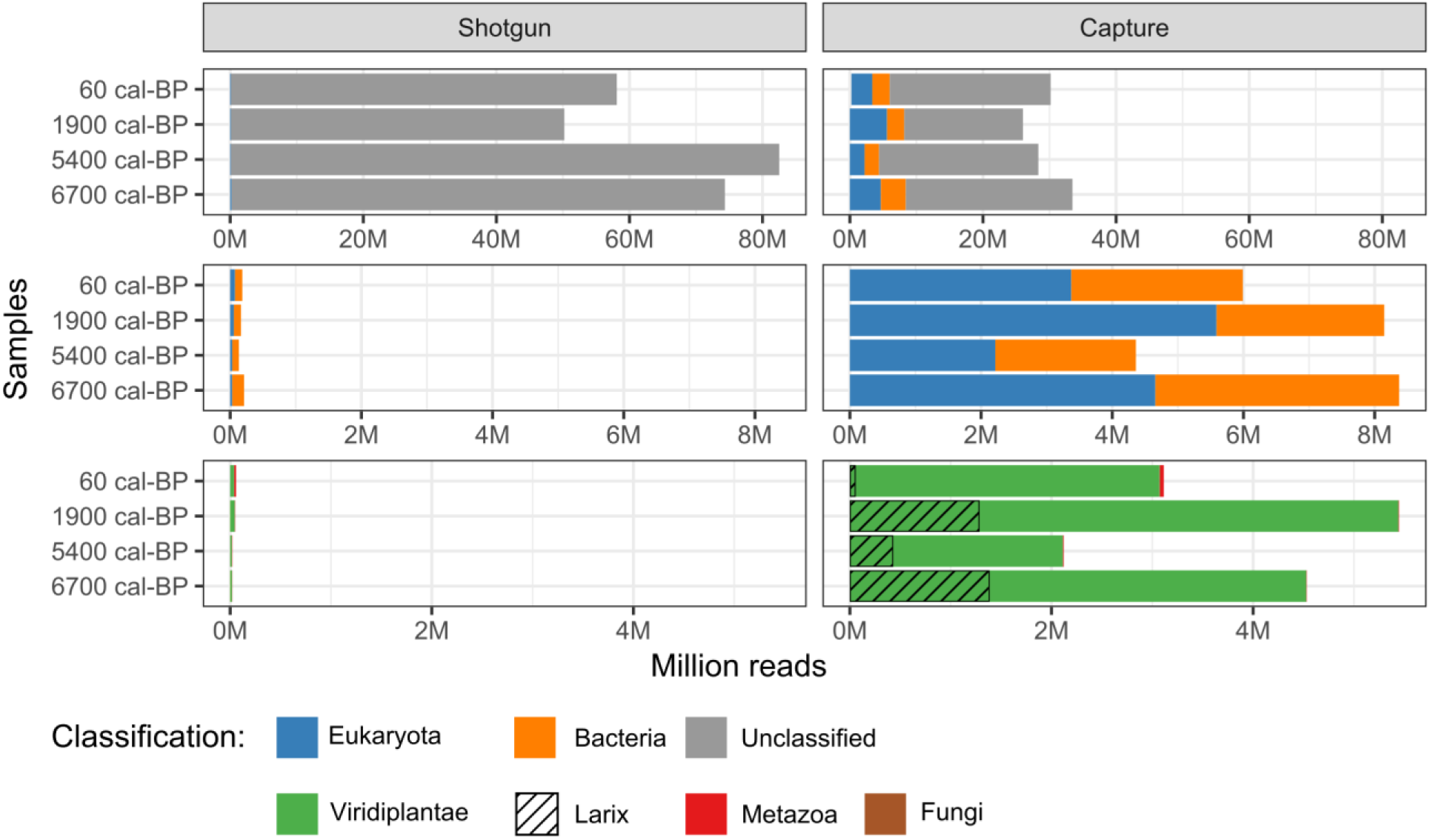
Sequence reads classified with kraken2 at high confidence against NCBI non-redundant nucleotide database (nt). Left: shotgun dataset, right: hybridization capture dataset. Upper graphs: classification at domain level. Middle graphs: as above with unclassified reads excluded. Lower graphs: classification of Eukaryota at kingdom level and classified as *Larix*.

The sequencing of the four hybridization capture samples resulted in approximately 192 M paired-end read counts with a per-sample read count of 42–50 M. Sequencing of the extraction blank and the library blank resulted in 4.3 and 5.7 M reads, respectively. After trimming and quality filtering, 66% of reads were kept from the samples (126 M reads) and 0.1% of reads were kept from the blanks (27 k reads). About 91% of the reads of each sample had overlapping reads and were merged. Classification with kraken2 using the nt database could classify 28% of the quality filtered reads. Of the classified reads, 31% were classified as Bacteria (11 M reads) and 44% were classified as Eukaryota (15.8 M reads). Of the latter, 15.2 M reads correspond to Viridiplantae and 3 M reads to *Larix* (Figure 1, Supplementary Table S1). A comparison of the shotgun and capture datasets shows that 46.6 to 155.8-fold more reads were assigned to Eukaryota in the capture dataset. On the taxonomic level of Viridiplantae enrichment ranged from 77.8 to 236.9-fold enrichment of captured data in respect to shogun data. The number of *Larix*-classified reads per sample corresponds to an increase of around 800 to 1160-fold compared to the shotgun data (Supplementary Table S2).

### Ancient DNA authenticity

A mapDamage analysis (Jónsson et al., 2013) was applied to the alignment files of *Larix-*classified reads aligned to the *L. gmelinii* chloroplast genome. The overlapping merged reads for the three ancient samples (from 1900, 5400 and 6700 cal-BP) show a clear increase of C to T substitutions at both ends with a greater pronunciation at the 5’ ends. A clear increase of substitution rate with age is visible (Supplementary Figure S4). The unmerged paired-end reads show comparable C to T substitution rates for the forward reads at the 5’ ends and for the reverse reads at the 3’ ends. The youngest sample (60 cal-BP) does not show any clear pattern of substitution rates for merged or unmerged paired-end reads. The length of sequencing reads was between 50 bp and 340 bp for all samples (Supplementary Figure S4 and S5).

### Retrieval of the *Larix* chloroplast genome

Sequence reads classified as *Larix* using a custom chloroplast database, were mapped to the *L. gmelinii* reference genome. In the hybridization capture experiment this resulted in a near-complete retrieval of the *Larix* chloroplast genomes for most samples. The coverage of the complete chloroplast genome declined from the oldest to the most recent sample. At a minimum of 1-fold coverage 91.4% of sample “6700 cal-BP” is covered, 80.3% of sample “5400 cal-BP”, 85.1% of sample “1900 cal-BP” and 14.3% of sample “60 cal-BP”. The mean coverage of the samples is 21.0x ± 14.2x (6700 cal-BP), 5.0x ± 4.5x (5400 cal-BP), 6.6x ± 5.9x (1900 cal-BP) and 0.3x ± 0.7x (60 cal-BP). Comparing the alignments of shotgun and capture reads, the enrichment ranged from 6.4 to 16.2-fold (Supplementary Table S2).

In the alignment of reads against the *L. gmelinii* reference chloroplast genome, the coverage is not equal across the different annotated regions. When aligning the *Larix-*classified reads, the coverage is highest for inverted repeats and lowest for ribosomal RNA (Figure 2, dark shaded colours). In the same dataset, the coverage is, on average, higher for intergenic regions, pseudogenes and conserved open reading frames (ORFs), than for protein coding genes. When aligning the complete (quality controlled) reads against the same reference, the coverages of the different annotated regions show a different pattern: highest coverage is at the ribosomal RNA, followed by the photosystem complex coding region and the inverted repeats.

**Figure 2.**
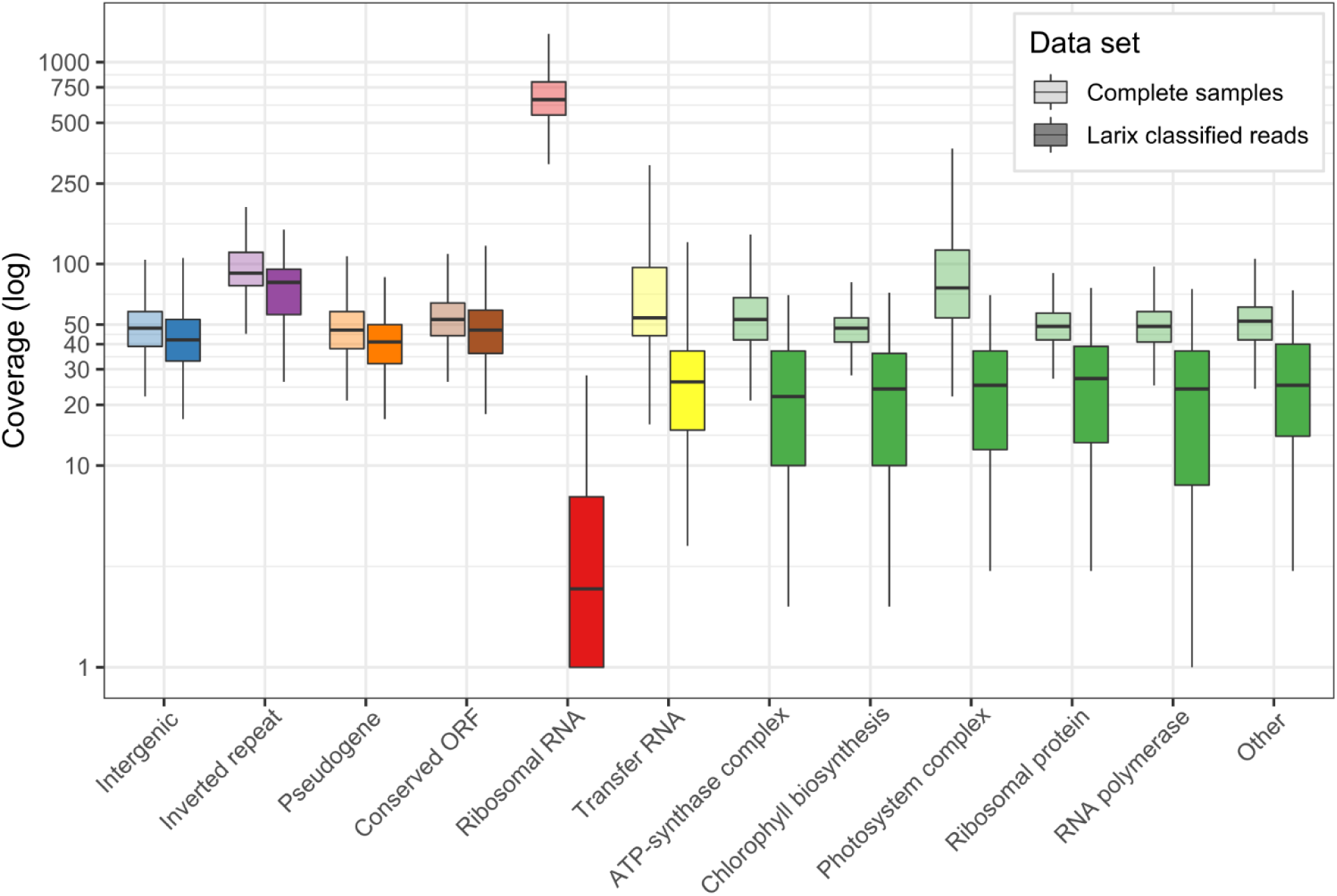
Read coverage of all four samples in the alignment of the capture dataset against the *L. gmelinii* chloroplast genome. Coverage is shown according to the functional annotation. Light colour shades: complete quality filtered read sets of the capture dataset; dark colour shades: *Larix*-classified reads of the capture dataset. Green shades: protein-coding genes. ORF: open reading frame of unknown function. Outliers not shown.

Considering the 294 sites which differ between the two reference genomes of *L. gmelinii* and *L. sibirica*, most of the reads carry *L. gmelinii* specific variations, but in all samples reads carrying the *L. sibirica* variants can also be found (Figure 3). Almost no reads carry neither of the two species-specific variations (“other”). Between 53% and 1% of the positions contained reads which were classified as *L. sibirica* variants, with the highest number detected at 6700 cal-BP and the lowest number detected at 60 cal-BP (Supplementary Table S3). The ratio of *L. sibirica* variants over all positions and reads varied from 5.17% (6700 cal-BP) to 1.84% (5400 cal-BP). Most of the variation between the two *Larix* species lie in the intergenic region and in conserved open reading frames (ORFs) of unknown function (Figure 3).

**Figure 3.**
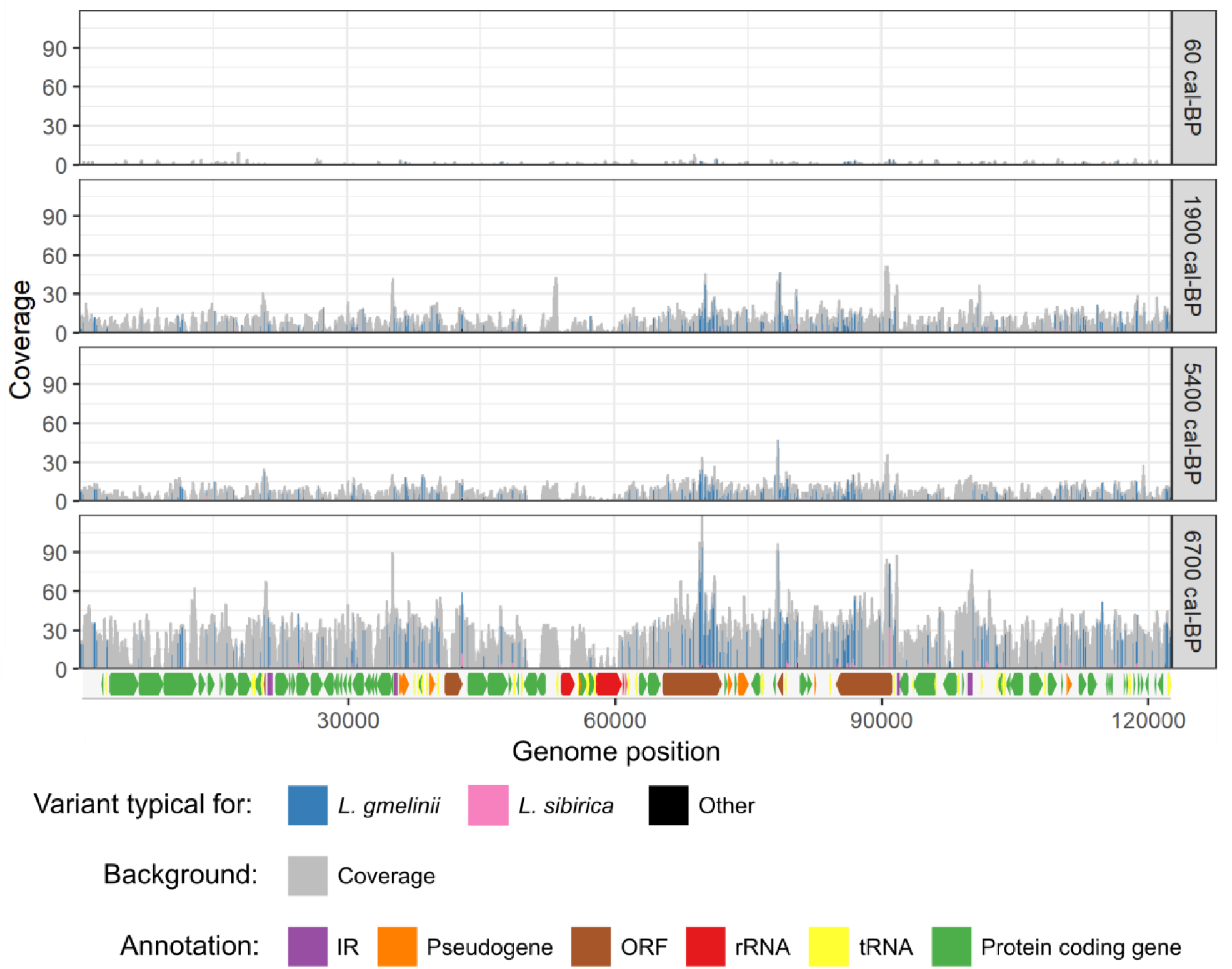
Alignment of *Larix* classified reads from hybridization capture dataset against the *Larix* chloroplast reference genome. The coverage per position is depicted in grey. For the 294 sites variable between the *L. gmelinii* and *L. sibirica* chloroplast genomes, colour indicates how many reads correspond to the variation found in each of the two species or if a read contained a variation found in neither of the two species (“other”). Key for the coloured arrows annotating the functional groups. IR: inverted repeat, ORF: open reading frame of unknown function, rRNA: ribosomal RNA, tRNA: transfer RNA.

## Discussion

Ancient DNA from lake sediments constitutes a valuable resource to investigate the response of populations to past environmental changes. Previous studies using metabarcoding or shotgun sequencing have not yet explored the full potential of this resource. Here, we applied shotgun sequencing and hybridization capture using PCR-generated baits of the *Larix* chloroplast genome, to retrieve nearly complete chloroplast genomes and study past changes in the population history of larches in northern Siberia.

### Taxonomic classification - majority of shotgun data could not be assigned

In the shotgun dataset, only 0.3% of quality filtered reads could be classified against the nt database. Considering that fewer than 15,000 organisms of the 10 to 15 million of eukaryotic and perhaps trillions of bacterial and archaeal species have currently been completely or partially sequenced (Lewin et al., 2018), this is a comparatively large fraction. However, other studies have reported higher rates of assigned sequences. Slon et al. (2017), assigned more than 10% of their shotgun reads that derived from cave sediments and Ahmed et al. (2018) reported 16% of assigned sequences from ancient lake sediments and 70% of hits over multiple taxa. On the other hand, Pedersen et al. (2016) assigned 0.3% of the quality filtered reads obtained from ancient lake sediments, and Parducci et al. (2019) reported a read assignment of 3.4% of *sed*aDNA shotgun sequencing. Different sample material, wet lab processing and sequence databases used for assignment make it difficult to properly compare the assignment rate. In addition to these differences, the applied bioinformatic approach has an impact on the results (Harbert, 2018). Here we used kraken2 (Wood, Lu, & Langmead, 2019), a new version of kraken, which is a particularly conservative tool compared to others, reporting less false-positive but also less true-positive hits than others tools, even with default values (Harbert, 2018). We used it with a high confidence threshold as we found it gives the best results in terms of vegetation composition based on our knowledge of the vegetation history (Epp et al., 2018), but with the consequence of very low overall assignment rates. Indeed, when we use the default values of kraken2, we could assign 10–16% of the reads. However, more lenient classification causes a reinforcement of the database bias: few deeply sequenced taxa are more likely to be assigned than the majority of shallowly or fragmentarily sequenced taxa (Parducci et al., 2017).

### Capture success - Viridiplantae reads increased by orders of magnitude

Hybridization capture resulted in an increase of taxonomically classified reads by orders of magnitude – especially with respect to the ratio of unclassified reads but also in total. These results show for the first time that DNA capturing of whole chloroplast genomes is effective even for DNA libraries that constitute a high diversity and low on-target rates such as DNA from ancient lake sediments. Along with *Larix* many other plant taxa were enriched, as is shown by the high number of Viridiplantae classified reads (Figure 1). This can be explained by: (1) chloroplast genomes of land plants and green algae sharing essentially the same set of protein-coding genes and ribosomal RNAs and differing mainly in the presence/absence of introns and repeats (B. R. Green, 2011), and (2) DNA libraries built from lake sediments contain a very high sequence divergence, higher than those from pooled individuals, which in the past were used to measure the capability of capturing highly divergent reads (Paijmans, Fickel, Courtiol, Hofreiter, & Förster, 2016; Peñalba et al., 2014). We hypothesize that the captured reads classified as other plants share, to some extent, sequence similarities with conserved regions like genes or ribosomal RNA.

The comparison of captured *Larix-*classified reads versus the full capture samples gives a first hint of this: if we align the complete dataset and the subset with the same relaxed parameters to the *Larix* chloroplast genome, the coverages are unequally distributed between the different functional regions. The conserved protein coding genes, especially genes coding for the photosystem complex, transfer RNA and ribosomal RNA, have coverages significantly higher in the complete dataset than in the *Larix-*classified dataset. In the case of ribosomal RNA the difference is even orders of magnitude higher (Figure 2). This shows that sequences from species other than *Larix* can align to these especially conserved regions. Several studies already demonstrate enrichment of diverged sequences using hybridization capture. Peñalba et al. (2014) show the applicability of hybridization capture to analyse taxa from 13 families with up to 27% sequence divergence, by targeting conserved gene regions. On ancient material, Paijmans et al. (2016) captured mitochondrial sequences with up to 20 million years divergence time with up to 25% sequence dissimilarity. With a modified relaxed double capture method, Li et al. (2013) demonstrate a successful enrichment of nuclear genes across distantly related species with up to 300 million years divergence time (with up to 40% sequence dissimilarity). All these studies demonstrate that hybridization capture can be used to study a wide range of phylogenetically divergent taxa, although they all note a decrease in coverage with increasing evolutionary distance.

### Near-complete retrieval of *Larix* chloroplast genomes

The number of reads classified as the genus *Larix*, using the nt database increased 800 to 1600-fold from shotgun to hybridization capture dataset. However, in all samples there was a high level of PCR duplicates due to the high number of amplifications after capturing and possibly also due to oversequencing of reads. Future projects could reduce the number of PCR cycles and increase the number of samples pooled together to reduce duplicates.

The deduplication of reads prior to mapping against the reference genome reduced the enrichment to 6–16-fold. This is in the range of results from enrichment studies performed on bone material (Avila-Arcos 2015, Carpenter 2013), although higher enrichment ranges have also been reported (Mohandesan 2017). Here, most likely the sample material plays a role, as the lake sediments contained a low content of the target larch chloroplast DNA together with a high sequence diversity. Nevertheless, the target enrichment rate could potentially be increased, for example, by capturing at higher temperatures or using a touch-down approach or by using RNA instead of DNA for baits (Carpenter et al., 2013; Paijmans et al., 2016; Peñalba et al., 2014).

In the capture dataset, near-complete chloroplast sequences could be retrieved from the three oldest samples. But even the sample with the highest coverage, i.e. the oldest from 6700 cal-BP, contained gaps in the alignment (91% covered at 1-fold coverage). When comparing the coverage of different functional groups of the genome (Figure 2), we see the lowest coverage for ribosomal RNA, followed by protein coding genes. This is due to the bioinformatics approach of extracting reads as classified to the lowest common ancestor in kraken2. As these coding regions are highly conserved across taxa (B. R. Green, 2011), the short reads can also be attributed to other organisms and are consequently classified to a higher taxonomic rank than *Larix.* Accordingly, when assigning the full, not taxonomically binned samples in the same manner to functional groups, this trend reverses.

Analysis of DNA damage patterns revealed C to T substitution rates typical for ancient DNA (Supplementary Figures S4, S5). Typical for the preparation of single-stranded libraries, these substitutions could be observed both at the 3’ and the 5’ end of the molecules (Gansauge & Meyer, 2013). C to T substitution rates increased with sample age, in line with previous observation (Pedersen et al., 2016; Sawyer, Krause, Guschanski, Savolainen, & Pääbo, 2012). Read lengths ranged from 40 bp to 340 bp, showing the short fragment length typical for ancient DNA (Green et al., 2008).

### *L. sibirica* variants present over time

When comparing the ancient reads to chloroplast reference genomes from *L. gmelinii* and *L. sibirica*, the great majority of reads carry the *L. gmelinii* variants with a low frequency of *L. sibirica* variants in all four samples. In contrast, the analysis of one mitochondrial marker derived from the same core by Epp *et al*. (2018), showed a mixture of both species, with relatively high rates of *L. sibirica* – except for the most recent sample, which showed clear dominance of the *L. gmelinii* mitotype – pointing to a co-occurrence of both species across most of the sediment core. In the genus *Larix*, chloroplasts are predominantly inherited paternally (Szmidt, Aldén, & Hällgren, 1987) whereas mitochondrial DNA is inherited maternally (DeVerno, Charest, & Bonen, 1993), a phenomenon which has been reported for almost all members of the conifers (Neale & Wheeler, 2019). This bi-parental inheritance results in different rates of gene flow and subsequently asymmetric introgression patterns (Du, Petit, & Liu, 2009; Petit et al., 2005). Simulations (Currat, Ruedi, Petit, & Excoffier, 2008) and molecular studies on a range of Pinaceae (Du et al., 2011, 2009; Godbout, Yeh, & Bousquet, 2012) showed that the seed-transmitted mitochondria which experience little gene flow introgress more rapidly than the pollen dispersed chloroplasts which experience high gene flow. A second finding of these studies is that introgression occurs asymmetrically from the resident species into the invading species. An expected result of introgression is therefore a population carrying mitotypes of the former local species and chlorotypes of the invaded species.

In the case of the population history of *L. sibirica* and *L. gmelinii* in their contact zone, Semerikov et al. (2013) found evidence for the asymmetric introgression of *L. sibirica* mitotypes in a population carrying only *L. gmelinii* chlorotypes, confirming the natural invasion of *L. gmelinii* into the range of *L. sibirica.* Here we corroborate these findings with a distinct discrepancy between relatively high rates of *L. sibirica* mitotypes as reported before (Epp et al., 2018) and low rates of *L. sibirica* in the chloroplast reads found in this study.

This points to an invasion of *L. gmelinii* in a former population of *L. sibirica* prior to the date of our oldest sample (6700 cal-BP). Further evidence in support of this scenario is found in the results from a lake sediment core 250 kilometres south-west of the study site (Epp et al., 2018), where samples reaching back to 9300 cal-BP show exclusively *L. sibirica* mitotypes, before they were gradually replaced by the *L. gmelinii* mitotype. Our study shows that by capturing for the complete chloroplast genome, we achieve a high resolution and can detect species specific variants even at low frequencies. Further studies should also include mitochondrial sequences in the target enrichment to collect data from several markers or potentially the complete mitochondrial genome. By combining the two organelle genomes in a hybridization capture experiment it would be possible to study hybridization and introgression events in detail, which would help to deepen our understanding of population dynamics over long time scales.

### Larch forest decline over the last 7000 years

When looking at the overall retrieval of *Larix* reads among the samples, most reads could be recovered from the oldest sample, of around 6700 cal-BP and the least number of reads in the most recent sample. This is in line with the findings of Klemm *et al*. (2016) and Epp *et al*. (2018), who describe a general vegetation turnover from larch forest to an open tundra with only sparse *Larix* stands during the last 7000 years at this site. This gradual change of vegetation in response to late-Holocene climate deterioration has also been observed by various studies (Andreev et al., 2004; MacDonald, Gervais, Snyder, Tarasov, & Borisova, 2000; MacDonald, Kremenetski, & Beilman, 2008; Pisaric, MacDonald, Velichko, & Cwynar, 2001) and is in correspondence with the reconstructed global cooling trend of the middle to late Holocene (Marcott, Shakun, Clark, & Mix, 2013).

## Conclusion

Siberian larch forest covers vast areas of northern Asia with *Larix* as the only tree-forming species. Lake sediments containing ancient DNA constitute an archive to answer the question of how larch forests respond to changing climate, but the low amount of target DNA in combination with a high degree of sequence divergence make them challenging material to study the population dynamics of a specific species. Here we have shown the success of hybridization capture of near-complete chloroplast genomes from 6700-year old lake sediments originating from northern Siberia. Shotgun sequencing of *sed*aDNA prior to enrichment showed that, depending on the cautiousness of the bioinformatic approach, only very low rates of reads can be securely assigned to taxa even at the domain level. By using PCR-generated baits covering the whole chloroplast of *Larix* for hybridization capture we could achieve increases by several orders of magnitude of assignable reads. The enrichment of *Larix* reads was most distinct, but plant DNA in general was also enriched. With ancient DNA from lake sediments, hybridization capture thus offers the potential of not only analysing the target species in depth, but also of studying the taxonomic diversity of the sample in a similar way to traditional molecular barcoding approaches. The method is more costly than the metabarcoding approach but has several advantages: no restriction to specific fragment length or by primer binding sites, avoidance of PCR introduced bias and the possibility of authentication of ancient DNA. Future studies focusing on plant biodiversity changes could focus on conserved coding regions of a set of diverged species to capture a more complete picture of the past vegetation. The analysed *Larix* reads confirm a general larch forest decline over the last 6700 years. Low rates of *L. sibirica* variants in proportion to *L. gmelinii* variants in the chloroplast could point to an invasion of *L. gmelinii* into *L. sibirica* populations before 6700 years ago. This study represents the first demonstration of hybridization capture from ancient DNA derived from lake sediments. Our results open the way for large scale palaeogenomic analyses of ancient population dynamics using lake sediment cores.

## Acknowledgements

We thank our Russian and German colleagues who helped in fieldwork in 2011 to obtain the samples. Nick Mewes is highly acknowledged for assistance in the laboratory. We also thank Cathy Jenks for English language proof reading. This project has received funding from the European Research Council (ERC) under the European Union’s Horizon 2020 research and innovation programme (grant agreement no. 772852, ERC consolidator grant “Glacial Legacy”) and the Initiative and Networking fund of the Helmholtz Association. LSE was supported by the German Research Foundation through grant EP98/2-1.

## Data Accessibility

The Illumina sequence data are submitted to the European Nucleotide Archive under project number PRJEB35838, accession numbers ERS4197088 - ERS4197099 (*Note: data will be released upon acceptance of the manuscript*).

## Author Contributions

LSE and UH designed the study; LS conducted library preparation and hybridization capture, analysed the data with the help of NB and HZ; LSE designed and supervised the preparation of the long-range PCR products; UH, NB, LE, KS and HZ supervised the study; LS wrote the manuscript under supervision of UH. All co-authors commented on a first version of the text.

