## Supplemental Information for "Hybridization capture of larch (*Larix* Mill) chloroplast genomes from sedimentary ancient DNA reveals past changes of Siberian forests"

*Table S1 Number of reads sequenced for shotgun sequencing and hybridization capture sequencing*

| Shotgun | 6700 cal-BP | 5400 cal-BP | 1900 cal-BP | 60 cal-BP | Ext. blank | Lib. blank |
|---|---|---|---|---|---|---|
| Raw reads | 145,972,964 | 112,220,286 | 78,650,028 | 87,627,075 | 10,370,556 | 14,264,796 |
| QC reads | 74,395,179 | 82,578,790 | 50,280,976 | 58,194,020 | 23,014 | 15,692 |
| Merged reads | 59,402,747 | 70,277,127 | 40,827,070 | 48,646,742 | 19,117 | 10,662 |
| Unclassified | 74,143,837 | 82,423,133 | 50,093,108 | 57,980,345 | 20,172 | 12,753 |
| Root | 251,342 | 155,657 | 187,868 | 213,675 | 2,842 | 2,939 |
| Archaea | 4,511 | 2,057 | 2,087 | 1,099 | - | - |
| Bacteria | 182,235 | 101,074 | 110,087 | 112,852 | 375 | 322 |
| Viruses | 98 | 938 | 60 | 107 | 3 | 2 |
| Eukaryota | 29,879 | 31,885 | 55,021 | 72,380 | 1,838 | 1,901 |
| Fungi | 494 | 1,273 | 898 | 961 | 9 | 10 |
| Metazoa | 2,011 | 3,530 | 3,004 | 17,940 | 1,797 | 1,874 |
| Viridiplantae | 19,115 | 17,270 | 46,359 | 39,477 | 30 | 16 |
| Larix | 1,196 | 495 | 1,150 | 45 | - | - |
| **Capture** | **6700 cal-BP** | **5400 cal-BP** | **1900 cal-BP** | **60 cal-BP** | **Ext. blank** | **Lib. blank** |
| Raw reads | 50,743,540 | 47,562,965 | 42,857,744 | 50,564,822 | 4,309,730 | 5,796,937 |
| QC reads | 36,512,567 | 30,066,323 | 27,832,438 | 32,166,404 | 19,898 | 6,788 |
| Merged reads | 33,115,641 | 27,802,780 | 25,304,175 | 29,301,513 | 19,898 | 6,788 |
| Unclassified | 25,071,453 | 23,956,488 | 17,858,027 | 24,178,726 | 8,885 | 3,156 |
| Root | 11,441,113 | 6,109,835 | 9,974,411 | 7,987,678 | 11,013 | 3,632 |
| Archaea | 41,891 | 16,271 | 21,801 | 11,507 | - | - |
| Bacteria | 3,718,173 | 2,142,634 | 2,558,563 | 2,616,197 | 1,641 | 772 |
| Viruses | 730 | 2,821 | 266 | 463 | - | - |
| Eukaryota | 4,654,826 | 2,217,357 | 5,587,144 | 3,373,420 | 5,112 | 896 |
| Fungi | 1,270 | 4,101 | 3,213 | 9,220 | 38 | 24 |
| Metazoa | 3,993 | 6,600 | 4,561 | 34,006 | 795 | 868 |
| Viridiplantae | 4,527,757 | 2,111,186 | 5,440,761 | 3,070,656 | 4,274 | 4 |
| Larix | 1,385,860 | 429,979 | 1,286,646 | 57,707 | 4 | 1 |

Ext. = extraction; Lib. = library; QC = quality control passed

*Table S2 Number of* Larix-*classified reads mapped to* L. gmelinii *chloroplast genome*

| | Capture | Shotgun | Enrichment |
|---|---|---|---|
| Merged reads | | | |
| 6700 cal-BP | 13906 | 1393 | 9.98x |
| 5400 cal-BP | 4125 | 478 | 8.63x |
| 1900 cal-BP | 3918 | 1149 | 3.41x |
| 60 cal-BP | 156 | 40 | 3.90x |
| Unmerged reads | | | |
| 6700 cal-BP | 10503 | 111 | 94.62x |
| 5400 cal-BP | 2175 | 29 | 75.00x |
| 1900 cal-BP | 4357 | 138 | 31.57x |
| 60 cal-BP | 201 | 7 | 28.71x |
| Sum of merged & unmerged | | | |
| 6700 cal-BP | 24409 | 1504 | 16.23x |
| 5400 cal-BP | 6300 | 507 | 12.43x |
| 1900 cal-BP | 8275 | 1287 | 6.43x |
| 60 cal-BP | 357 | 47 | 7.60x |

*Table S3 Number of reads classified as* L. gemlinii *or* L. sibirica *in the alignment of Larix-classified reads against the* L. gmelinii *chloroplast genome at the 294 variable sites*

| | Sample | *L. gmelinii* | *L. sibirica* | other |
|---|---|---|---|---|
| No. of sites occurrence detected | 60 cal-BP | 59 | 3 | 4 |
| | 1900 cal-BP | 283 | 43 | 14 |
| | 5400 cal-BP | 278 | 29 | 9 |
| | 6800 cal -BP | 289 | 157 | 36 |
| Total number of reads per classification | 60 cal-BP | 116 | 5 | 5 |
| | 1900 cal-BP | 2958 | 84 | 20 |
| | 5400 cal-BP | 2125 | 40 | 11 |
| | 6800 cal -BP | 8301 | 455 | 48 |
| Percent of total reads | 60 cal-BP | 92.1 | 4.0 | 4.0 |
| | 1900 cal-BP | 96.6 | 2.7 | 0.7 |
| | 5400 cal-BP | 97.7 | 1.8 | 0.5 |
| | 6800 cal -BP | 94.3 | 5.2 | 0.5 |
| Percent of sites with occurrence detected | 60 cal-BP | 20.1 | 1.0 | 1.4 |
| | 1900 cal-BP | 96.3 | 14.6 | 4.8 |
| | 5400 cal-BP | 94.6 | 9.9 | 3.1 |
| | 6800 cal -BP | 98.3 | 53.4 | 12.2 |
| Median of total number of reads per classification | 60 cal-BP | 0 | 0 | 0 |
| | 1900 cal-BP | 9 | 0 | 0 |
| | 5400 cal-BP | 7 | 0 | 0 |
| | 6800 cal -BP | 27 | 1 | 0 |
| Mean number of reads total per classification | 60 cal-BP | 0.39 | 0.02 | 0.02 |
| | 1900 cal-BP | 10.06 | 0.29 | 0.07 |
| | 5400 cal-BP | 7.23 | 0.14 | 0.04 |
| | 6800 cal -BP | 28.23 | 1.55 | 0.16 |
| Standard deviation of total number of reads per classification | 60 cal-BP | 0.9 | 0.2 | 0.2 |
| | 1900 cal-BP | 7.8 | 0.8 | 0.3 |
| | 5400 cal-BP | 5.1 | 0.5 | 0.2 |
| | 6800 cal -BP | 13.6 | 2.6 | 0.5 |

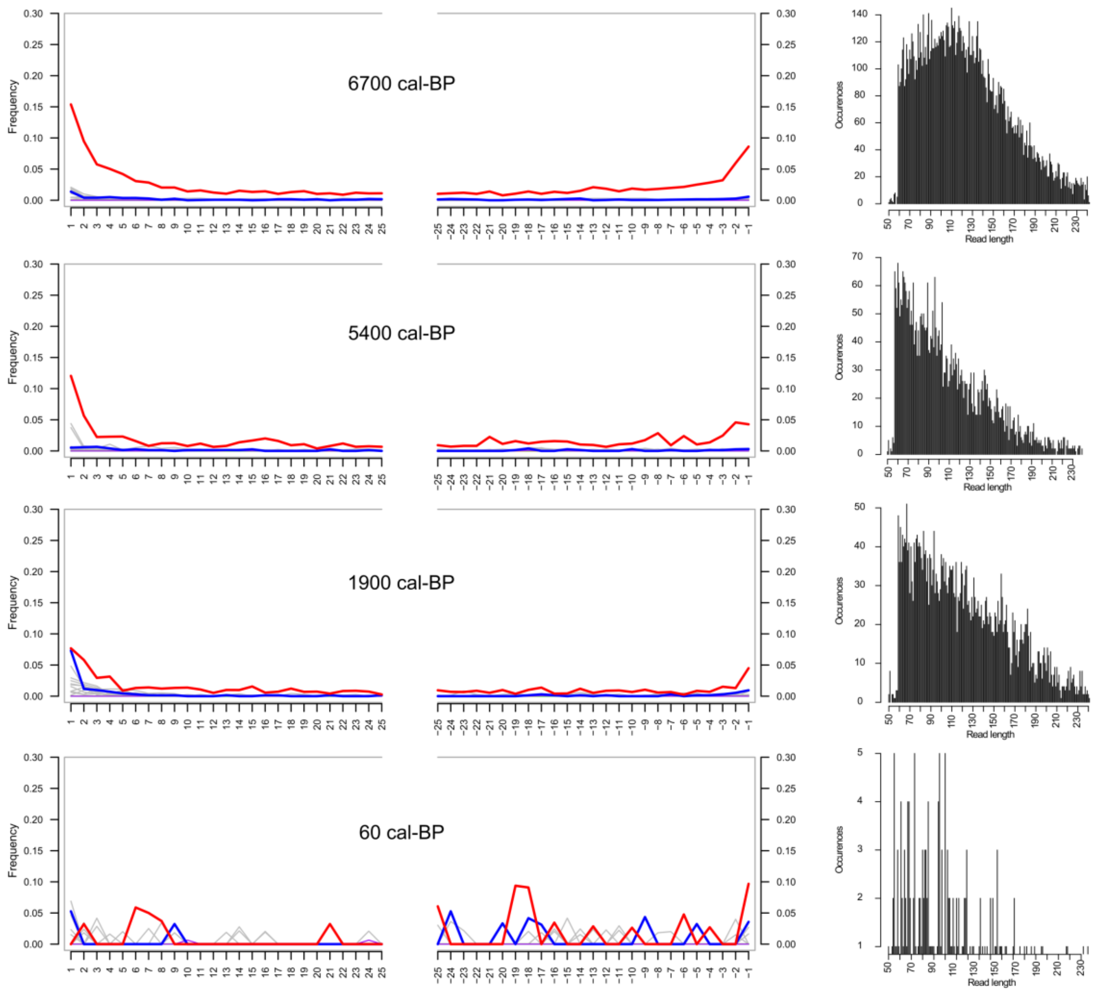


Figure S1 MapDamage plots for overlapping merged reads of the hybridization capture dataset aligned against the *Larix* chloroplast genome: Left: Misincorporation plots, red: C to T substitutions, blue: G to A substitutions, grey: all other substitutions. Right: Read length distributions.

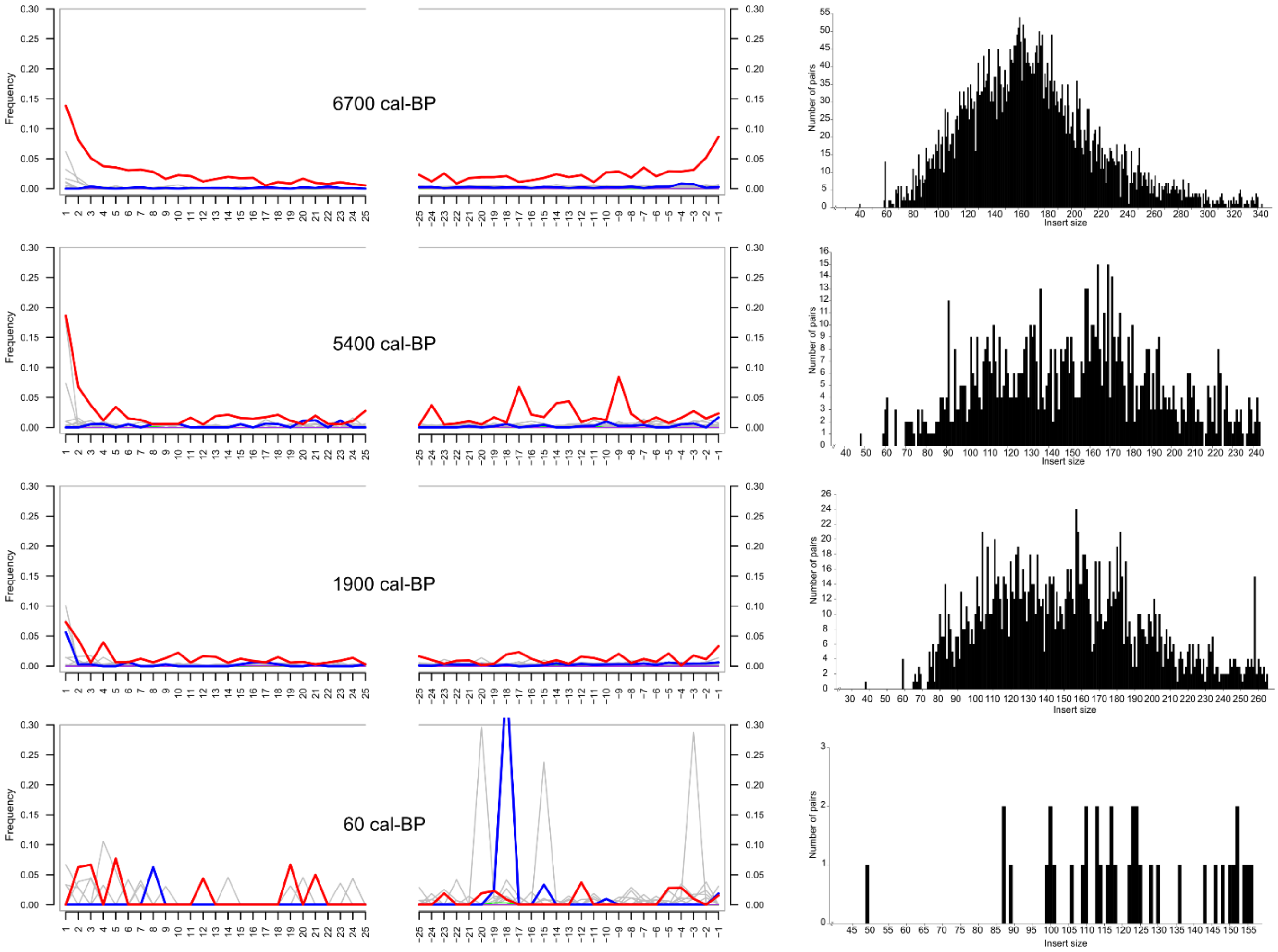


Figure S2 MapDamage plots for unmerged paired end reads of the hybridization capture dataset aligned against the *Larix* chloroplast genome: Left: Mis-incorporation plots, left site from analysis with only forward reads, right site from analysis with only reverse reads. Right: Insert size distribution for mapped paired-end reads. Adapted from geneious plot
